# Therapeutically exploring persister metabolism in bacteria

**DOI:** 10.1101/737320

**Authors:** Sayed G. Mohiuddin, Thuy Hoang, Adesola Saba, Prashant Karki, Mehmet A. Orman

## Abstract

Bacterial persisters are rare phenotypic variants that are temporarily tolerant to high concentrations of antibiotics. We have previously discovered that persisters are mostly derived from stationary-phase cells with high redox activities that are maintained by endogenous protein and RNA degradation. This intracellular degradation resulted in self-inflicted damage that transiently repressed the cellular functions targeted by antibiotics. Leveraging this knowledge, we developed an assay integrating a degradable fluorescent protein system and a small library, containing FDA-approved drugs and antibiotics, to detect chemicals that target persister metabolism. We identified several metabolic inhibitors, including anti-psychotic drugs, that can reduce *Escherichia coli* persistence. These chemical inhibitors also reduce *Pseudomonas aeruginosa* persistence, potentially verifying the existence of similar mechanisms in a medically relevant organism.

## INTRODUCTION

Conventional therapies for infectious diseases target the mechanisms that enable the rapid growth of bacterial cell populations. Although this can provide a clinical benefit, this benefit is usually short-lived for persistent and recurrent infections, and a large body of evidence suggests that small subpopulations of microbial cells invariably survive this initial selection pressure. One of the proposed mechanisms for this tolerance is via the establishment of a latent pool of persister cells (1). Persisters are an important health problem, because they are thought to underlie the propensity of recurrent infections to relapse (2–4) and serve as a reservoir from which drug-resistant mutants can emerge (5–8). Persisters exhibit a diverse range of proliferative, metabolic, and transcriptional activities. Whereas there are some variants that can grow in the presence of antibiotics, these are very rare and often survive the drug treatments by activating drug efflux systems (9) or bypassing the pathways targeted by the drugs (10). By contrast, the most abundant variant is the type I persisters, which do not grow in the presence of antibiotics and are largely formed by passage through the stationary phase before antibiotic treatments (11). Elucidating the formation mechanisms of these preexisting, nonproliferating type I persisters is of special interest; because, these variants are found among many bacterial species, are often multidrug tolerant, and their eradication is a huge challenge (1, 3, 4).

We previously showed that type I persisters mostly derive from stationary-phase cells with high redox activities that are maintained by endogenous protein and RNA degradation (12). We speculated that this intracellular degradation (i.e., self-digestion or autophagy) not only provides energy to bacterial cells in a non-nutritive environment, but also produces self-inflicted damage that renders the cells less fit for rapid resumption of growth. Inhibiting stationary-phase respiratory activities chemically (treatment with potassium cyanide or nitric oxide to suppress cellular respiration), environmentally (culturing under anaerobic conditions), or even genetically (genes encoding redox enzymes such as *ubiF, sucB, mdh, aceE, sdhC*, and *acnB*) reduced persister levels by preventing digestion of endogenous proteins and RNA, yielding cells that were more capable of translation and replication and thus susceptible to cell death when exposed to antibiotics (12, 13). This reduction in persister levels was not found to be associated with the inhibition of RNA and protein synthesis or elimination of reactive oxygen species (ROS) (12). These results also suggest persisters harbor ETC activities associated with bacterial cytochromes, oxidoreductases and PMF, which was supported by previous studies, where “aminoglycoside (AG) potentiation assays” were used (14–16). Our current study further provides strong support for the notion that persister metabolism is a rich source of novel antipersister strategies. Using a high-throughput screening approach and a small chemical library (Biolog Phenotype Arrays containing FDA-approved drugs and antibiotics) in the current study, we identified a subset of drugs that can reduce persistence in Gram-negative bacteria by targeting their metabolism.

## RESULTS

### Chlorpromazine pretreatment can reduce *E. coli* persistence

As effective sterilization methods for treating chronic and recurrent infections remain scarce, identifying novel targets, together with medicinally relevant inhibitors, is becoming an urgent priority to improve the therapies for these infections. Inhibition of respiration throughout stationary phase or deletion of genes encoding TCA and ETC enzymes was shown to delay intracellular degradation and persistence by reducing cellular metabolic activity (12). ETC reactions are powered by oxidizing/reducing equivalents and are essential for ATP generation by the proton motive force (PMF). If this is the case, we should be able to reduce persister formation in stationary phase by targeting a key component, i.e., ATP synthase, in this metabolic mechanism. Chlorpromazine, which is an FDA approved antidepressant drug that is effective, safe and listed as an essential medicine by the World Health Organization (17), was demonstrated to inhibit the catalytic complex of rotary nanomotor ATP synthase (F1-ATPase) in *E. coli* cells (18, 19). As expected, when we treat stationary-phase cultures with chlorpromazine (Fig. 1A), at a concentration that does not affect stationary-phase cell survival (Fig. S1), we were able to reduce the stationary-phase persistence (Fig. 1BC). We note that cells were not treated with antibiotics directly in stationary-phase cultures, as normal cells are intrinsically tolerant in these cultures (14, 16). The stationary-phase cells were first washed to remove the metabolic inhibitors, transferred to fresh medium, and then treated with antibiotics to stimulate non-persister cell killing. Pretreatment with chlorpromazine also reduced stationary phase metabolic activities by redox sensor green (RSG) dye (Figs. 1DE). RSG can readily penetrate bacteria and yield green fluorescence when reduced by bacterial reductases; hence, fluorescent signals produced by RSG correlate with cellular metabolic activities (Fig. S2). Overall, these results verify that bacterial metabolism is a rich source of novel antipersister strategies.

**Fig. 1.**
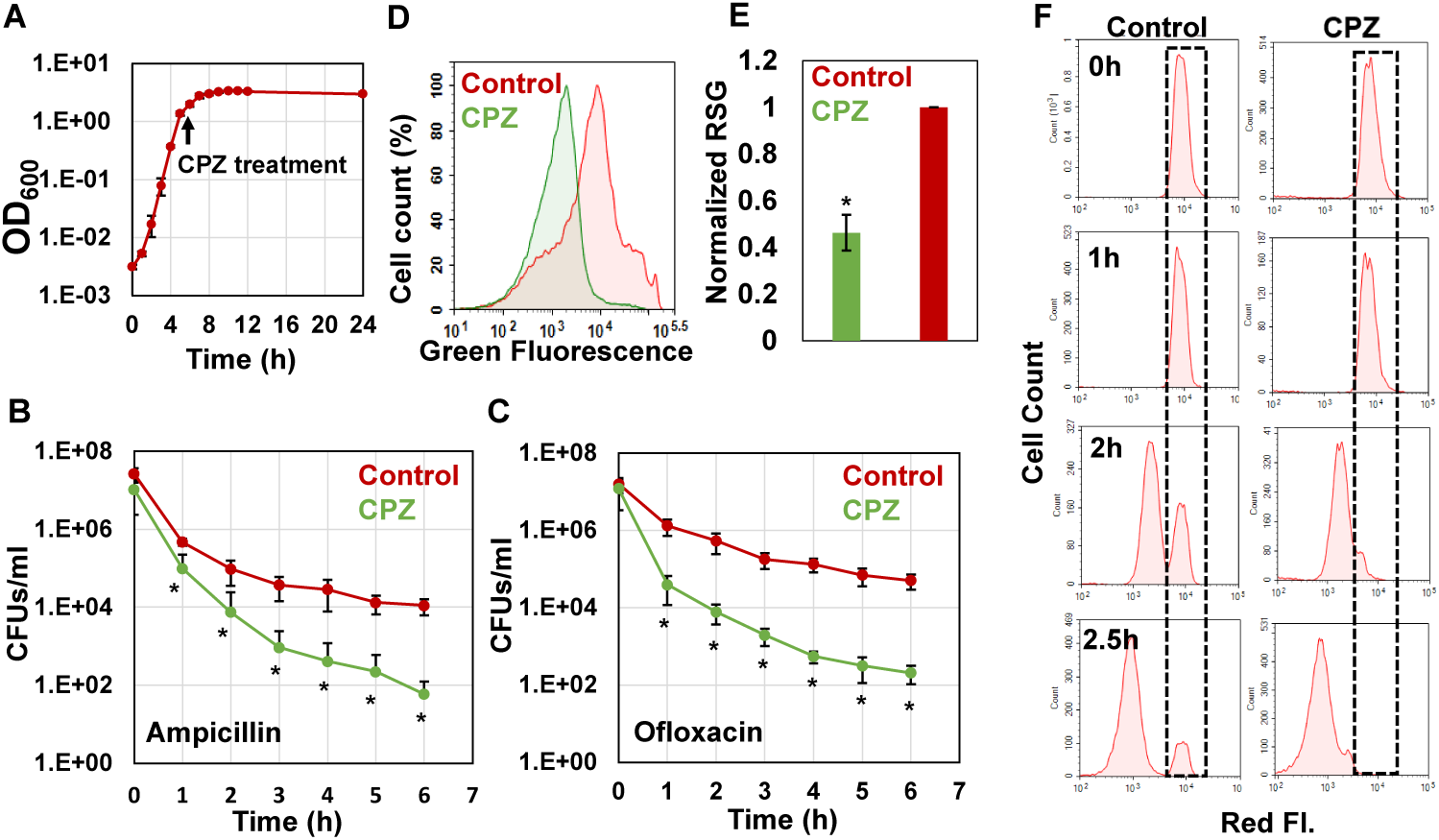
Chlorpromazine (CPZ) treatment reduced stationary-phase persistence, redox activities and non-growing cell formation. **(A)** CPZ treatment. Early-stationary-phase cells (t=5h) were treated with 0.25-mM CPZ or left untreated (control); cells in late-stationary phase (t=24h) were then washed to remove inhibitors to measure persister and non-growing cell levels as well as cellular redox activities. Cell growth was monitored with OD_600_ measurements. **(B-C)** Persister levels of CPZ-treated cultures. Untreated or CPZ-treated cells in late-stationary phase (t=24h) were washed to remove chemicals and resuspended in fresh media with antibiotics for persister quantitation (Number of biological replicates, N=6). **(D-E)** RSG staining of CPZ-treated or untreated late-stationary-phase cells. Untreated or CPZ-treated cells in late-stationary phase (t=24h) were washed to remove the chemicals, and resuspended in PBS to stain with RSG (N=6). **(F)** Non-growing cell levels in CPZ-treated cells. Early-stationary-phase cells (harboring an IPTG inducible mCherry expression cassette) were treated with 0.25-mM CPZ or left untreated (control) at t= 5h in the presence of IPTG; cells in late-stationary phase (t=24h) were washed to remove the chemicals and diluted in fresh media without inducer. Division at the single-cell level was monitored by flow cytometry during exponential-growth phase. A representative biological replicate is shown here. All 3 biological replicates consistently resulted in similar trends. *: Statistical significance between control (untreated) vs. CPZ treatment group at indicated time points or conditions (P<0.05, two-tailed t-tests with unequal variances).

### Chlorpromazine pretreatment reduced non-growing cell levels in stationary phase cultures

Although some persistent infections are associated with clinically apparent chronic symptoms, some cases are asymptomatic for a long period of time (e.g., a decade) and can develop clinically significant diseases at later times (20). The bacteria causing asymptomatic infections can be present within the host system in a nonreplicating or slowly replicating state (generally referred to as “viable but non-culturable” or VBNC state) and cannot be easily cultured in vitro (21, 22). We and others have shown that antibiotic-treated cultures have many more VBNC cells than persisters (∼2-log-fold more) (14, 23–25). Both persister and VBNC cells are stained as live, retain metabolic activity, and often appear as nongrowing during the antibiotic treatment (14). The only means to distinguish these subpopulations lies in the ability of persisters but not VBNC cells to recolonize in standard culture media in the absence of antibiotics. To determine whether the chlorpromazine pretreatment eliminate VBNC, we used our published method where we monitor cell proliferation via an inducible fluorescent protein (mCherry) expression cassette (12, 14, 26), in which mCherry-positive cells from late-stationary-phase cultures are inoculated in fresh medium in the absence of inducer (Fig. 1F, t=0). Flow cytometry reveals ongoing cell division as a dilution of mCherry, whereas the fluorescence levels are maintained in the nonproliferating subpopulation (Fig. 1F, WT at t=2.5 h). Although persisters were shown to be enriched in this subpopulation (26), most of these non-growing cells were identified as VBNC cells (26) (Fig. S3AB), which were not detected in the chlorpromazine treated cultures (Fig. 1F, Chlorpromazine at t=2.5h, and Fig. S3A). The reduction in both persister and VBNC cell levels in the chlorpromazine treated cultures points out these two phenotypes may be related. Consistent with the general notion in the field, it is possible that persistence may be a transitory phase leading to the VBNC state (22). Whether persistence contributes to the accumulation of VBNC cells due to the catabolism of intracellular components warrants further investigation.

### High-throughput screening detected chemical compounds that target *E. coli* metabolism and persistence

To directly measure protein degradation rates in stationary-phase cultures, we previously developed an assay using GFP that is linked to a short peptide degradation tag (11 amino acid residues), ssrA, to mark it for degradation by cellular proteases, mainly ClpAP and ClpXP (Lon, Tsp and FtsH are also known to target the ssrA sequence) (27–30). Although we note that self-digestion is a complex network orchestrated by many degradative enzymes (proteases, RNases and toxins), chlorpromazine treatment suppressed degradation of this tag in stationary-phase cultures (Fig. 2A, and Fig. S4A), potentially by reducing stationary-phase cell metabolic activities (Fig. 1DE). To test whether this straightforward system can identify additional antipersister therapeutics, we used a small library (Biolog Phenotype Arrays), containing antibiotics and other FDA approved drugs among ∼360 known chemical compounds in 96-well plate formats. Cells expressing ssrA-tagged GFP were transferred to the phenotype arrays without inducer at early-stationary phase, and cultured under the conditions studied here (Fig. 2B, see Materials and Methods). GFP levels were monitored using a plate reader, with cells cultured in the presence of the solvent serving as the negative controls, and those with chlorpromazine as a positive control. Our data verify that GFP in negative-control wells is degraded within 4 h (Fig. S4AB). The Z-factor (predicted by analysis of test plates with negative and positive controls, as described in Materials and Methods) was calculated to be 0.836, which indicates the robustness of our methodology (31). We employed a widely used Z-score method (calculated from the mean and the standard deviation of all measurements within the plate) (32) to determine initial hits. An absolute Z-score of ≥2 is the threshold for hit detection (32). Given that each plate contains four different concentrations for each compound (information on these concentrations not disclosed by the company), the initial hits were selected among the chemicals that successfully inhibited GFP degradation (Z-score≥2) with at least two different concentrations (Fig. 2C and Fig. S4C). As expected, chlorpromazine, which is one of the 360 chemical compounds tested, was identified as a positive hit, verifying that our method can detect potential metabolic inhibitors (Fig. 2C). To determine chemical inhibitors that specifically target persister metabolism, the identified hits were further analyzed in additional rounds of screening to determine concentrations that lead to complete inhibition of GFP degradation without affecting the stationary-phase-cell viability (Fig. 2D, and Fig. S5 and S6). We identified that CCCP, polymyxin B, poly-L-lysine, thioridazine, and trifluoperazine did not drastically affect the cell viability at the inhibitory concentrations for GFP degradation (Fig. S6), and four drugs, except CCCP, were able to reduce persistence (Fig. 2E and Fig. S7). Both thioridazine and trifluoperazine fall under the category of phenothiazine antipsychotic drugs, which are tricyclic compounds structurally similar to chlorpromazine. These drugs have been shown to reduce or inhibit NADH2-menaquinone-oxidoreductase and succinate dehydrogenase activities as well as altering NADH/NAD ratios (33–35), consistent with our RSG staining results provided in Fig. 3AB. We observe similar reduction in stationary phase cellular redox activities after polymyxin B and poly-L-lysine treatments (Fig. 3AB). These cationic peptides were shown to electrostatically bind to bacterial cells that leads to possible disruption of the bacterial membranes and membrane potential (36, 37), which explains the observed reduction in bacterial redox activities (Fig. 3AB). Treating the stationary phase cells with these four chemicals further reduces VBNC formation (Fig. 3C and Fig. S8), consistent with the results obtained from chlorpromazine treatments. Overall, these results strongly support that the identified drugs eliminate bacterial persistence by inhibiting stationary phase metabolism.

**Fig. 2.**
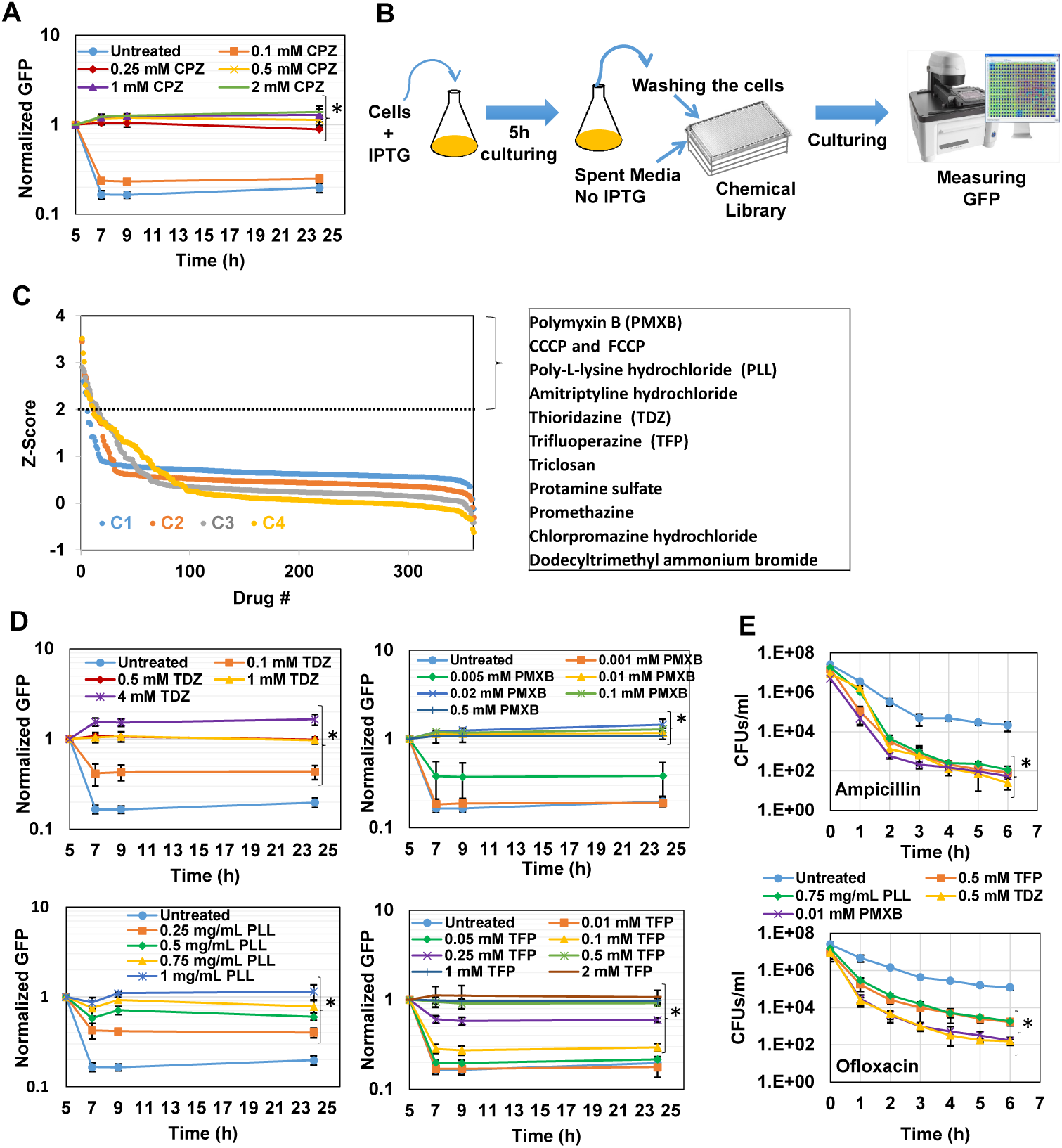
High-throughput drug screening detected chemical compounds that inhibit persistence. **(A)** Inhibition of GFP degradation with CPZ treatment at indicated concentrations. Cells expressing pQE-80L*gfp-ssrA* were grown to stationary phase (t=5h) in the presence of IPTG (inducer) and then re-suspended in a filter-sterilized spent medium (without inducer and obtained from the cultures grown under identical conditions) and immediately treated with CPZ to inhibit cell metabolism and protein degradation. Green fluorescence levels were measured and normalized to their initial levels (t=5h, before CPZ treatment) to determine GFP degradation. Background fluorescence was determined using cells with empty vectors (N=3). **(B)** High-throughput drug screening approach to identify chemical compounds that inhibit GFP degradation. Stationary-phase bacterial cells expressing ssrA-tagged GFP were re-suspended in spent medium, without inducer, transferred to 96-well PM plates containing the chemical library, covered with sterile, oxygen-permeable sealing membranes, and cultured in a shaker for 4h. GFP measurements taken at 4 h were normalized to those taken at 0 h (after transferring the cells to plates). **(C)** The Z-scores calculated for the chemical compounds at four different concentrations (C_4_> C_3_> C_2_> C_1_). Note that these concentrations were not disclosed by Biolog, Inc. The initial hits tabulated were selected among the chemicals that have Z-scores ≥2 with at least two different concentrations. **(D-E)** Inhibition of GFP degradation and persistence by the identified drugs. The selected hits were analyzed in depth at various concentrations to select the drugs that can reduce GFP degradation and persistence without affecting the *E. coli* cell viability. Cells were treated with these drugs at early stationary phase (t=5h) at indicated concentrations, and then, GFP measurements were performed at indicated time points. Persister assays were performed at late stationary phase (t=24h) (N=3). *: Statistical significance between drug-treated vs. untreated cultures at last three time points (P<0.05, two-tailed t-tests with unequal variances). CPZ: Chlorpromazine; PMXB: Polymyxin B; PLL: Poly-L-lysine; TDZ: Thioridazine; TFP: Trifluoperazine.

**Fig. 3.**
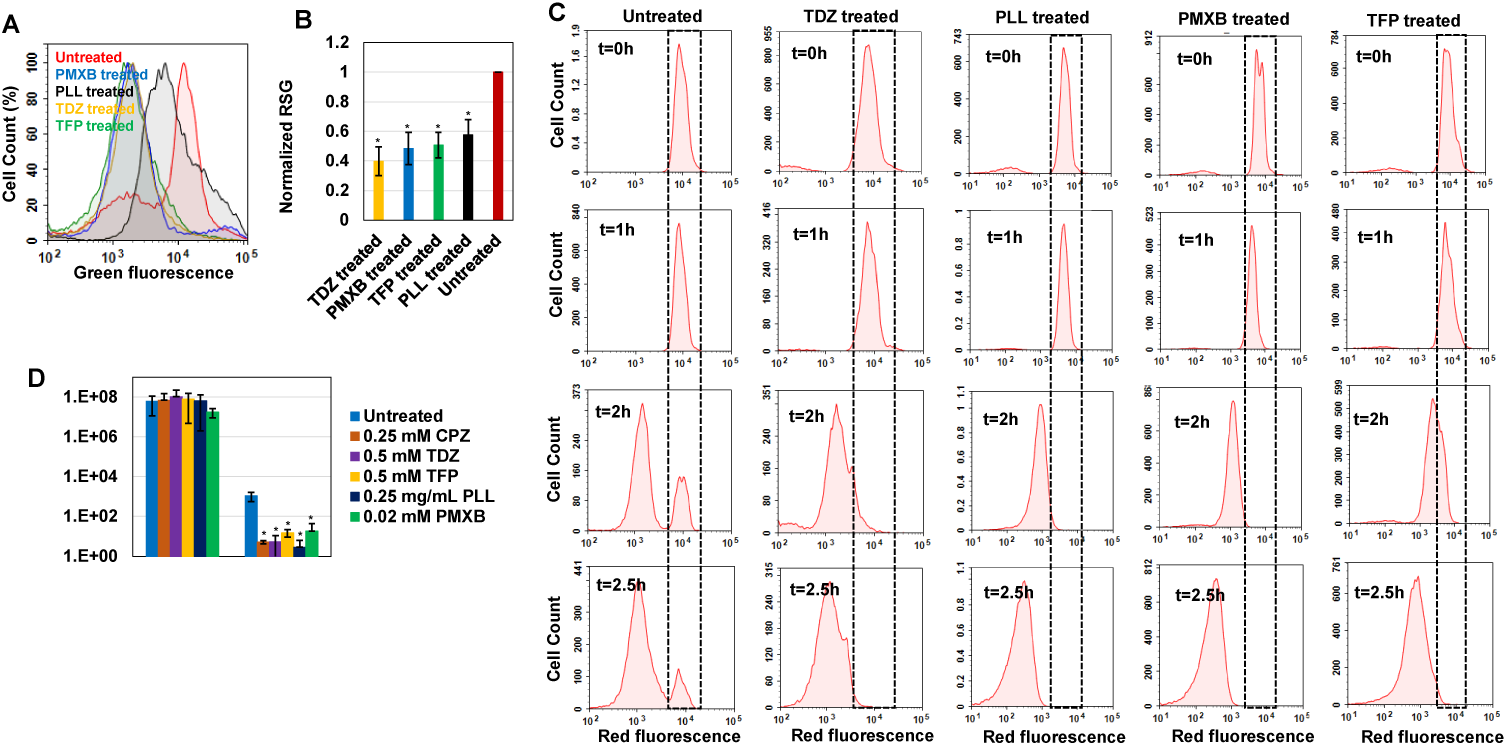
Drug treatments reduced stationary-phase redox activities and non-growing cell formation. **(A-B)** RSG staining of drug-treated or untreated late-stationary-phase *E. coli* cells. Cells were treated with the drugs at early stationary phase (t=5h), and RSG staining was performed at late-stationary phase (t=24h). Drug concentrations: 0.5 mM Thioridazine (TDZ); 0.75 mg/ml Poly-L-lysine (PLL); 0.01 mM Polymyxin B (PMXB); 0.5 mM Trifluoperazine (TFP) (N=6). **(C)** Non-growing cell levels in drug-treated *E. coli* cultures. A representative biological replicate is shown here. All 3 biological replicates consistently resulted in similar trends. Drug concentrations are the same as those provided in panels A-B. **(D)** Persister levels in *P. aeruginosa* cultures treated with the chemical hits. Early-stationary-phase cells (t=5h) were treated with the selected drugs or left untreated (control); cells in late stationary phase were then washed to remove inhibitors and re-suspended in fresh media with ofloxacin (effective for *P. aeruginosa*) (58) for persister assays. Cells were plated for CFU enumeration before and after the ofloxacin treatments to assess the effects of drugs on *P. aeruginosa* cell viability and persistence, respectively (N=6). *: Statistical significance between drug-treated vs. untreated cultures (P<0.05, two-tailed t-tests with unequal variances).

### The identified drugs can reduce *Pseudomonas aeruginosa* persistence

Our previous results indicate that persistence is facilitated by a self-digestion mediated metabolic futile cycle, wherein energy derived from catabolism is dissipated through continuous degradation of cellular components (12, 13). This process also introduces a self-inflected damage in the cells that transiently repressed the cellular functions targeted by antibiotics (12). The identification and characterization of the main components of this metabolic cycle may provide a global treatment approach as it can be an evolutionarily conserved process that may occur in many prokaryotes and eukaryotes and enable survival under stressful conditions (such as nutrient depletion, aging and overpopulation) via the recycling of essential energy molecules. When we similarly tested the identified chemicals on *Pseudomonas aeruginosa* (PAO1), we were able to substantially reduce *P. aeruginosa* persistence, suggesting the existence of similar mechanisms in other bacteria (Fig. 3D and Fig. S9). These results provide further clinical relevance for the identified drugs, since *P. aeruginosa* is involved in many hospital-related biofilm infections and the predominant cause of morbidity and mortality in cystic fibrosis patients with compromised immune systems (38–40).

## DISCUSSION

As antibiotics are most effective against growing bacteria, the resistance of persisters has been attributed to transient growth inhibition. Experimental evidence supporting this hypothesis was obtained in 2004 by Balaban and colleagues, who showed bacteria that failed to replicate prior to an ampicillin challenge also failed to lyse or grow during antibiotic treatment, but began replicating once the antibiotic was removed (11). This seminal study led to the model that persistence is a dormant phenotype, characterized by a depressed metabolism. However, recent evidence suggests persisters can harbor electron transport chain (ETC) activities associated with bacterial cytochromes and oxidoreductases (12). They can consume certain carbon sources to generate proton motive force (PMF) (14, 15), maintain high ATP levels (41), and drive the futile production and degradation of RNA, leading to energy generation and dissipation (42). Interestingly, most persister-related genes identified so far either directly or indirectly modulate cell metabolism.

Although metabolic processes and persistence in bacteria are known to be closely related, the specific mechanisms that link these remain unknown. Our previous results indicate that self-digestion may be this link (12). The role of metabolism is significant for bacteria, because they must produce large amounts of energy and biosynthetic precursors to meet the metabolic demands of their rapid growth. This results in a number of metabolic stresses, including nutrient starvation, hypoxia, and oxidative stress, which promote intracellular degradation/damage that may transiently repress the cellular functions targeted by antibiotics. Using transmission electron microscopy (TEM) and classic starvation conditions to create VBNC cells, Kim *et al*. showed that prolonged nutrient deprivation (7 weeks) results in cells that are spherical, have an empty cytosol (due intracellular degradation), and fail to resuscitate (43). Although nutrient deprivation initially increased persister levels in their experiments, continuous intracellular degradation eventually converted most of the cells to VBNCs. Persistence may, in fact, represent a transitory phase leading to the VBNC state and contribute to accumulation of VBNC cells due to intracellular degradation. Many persistence mechanisms identified so far involve stress-related responses, which generally induce, or are associated with, cellular self-digestion (44).

Although our previous results provided evidence that intracellular degradation transiently induces persistence (12), knowledge regarding what unique metabolic mechanisms are involved is lacking. Our current and previous results indicate that, despite their non-proliferating state, persister cells still exist in a metabolic steady state, where energy is continually produced and consumed (12–14). Our results further showed that targeting persister metabolism holds great potential for eradicating these dangerous phenotypes, as verified by the identified drugs (i.e., chlorpromazine, thioridazine, trifluoperazine, polymyxin B and poly-L-lysine), which are already known to target bacterial redox activities (45). Chlorpromazine (CPZ), thioridazine (TDZ), and trifluoperazine (TFP) are commonly known as first generation antipsychotic/neuroleptic drugs (45–47). Since they are the derivative of a heterocyclic phenothiazine, their mechanism of action is similar (45). The effectiveness of these drugs depends upon the ability to block dopamine receptors as the excessive dopamine is the main culprit of schizophrenia and other psychotic diseases (48). These drugs were also shown to have antimicrobial activities. In *Mycobacterium tuberculosis*, phenothiazines inhibit cellular respiration, leading to depletion of ATP as well as the reduction of NADH/NAD+ and menaquinol/menaquinone ratios (33–35). Because of their ability to inhibit bacterial efflux pumps, they were also shown to enhance the sensitivity of *Staphylococcus aureus* to beta-lactam antibiotics (49, 50).

Studies have shown that poly-l-lysine, which is a cationic polymer, can result in change of morphology in bacteria (37). In addition, treatment with poly-l-lysine raises the electric conductivity of the bacterial cells which leads to possible disruption of the cytoplasmic membrane (51). Similarly, polymyxins consist of a polypeptide cationic ring made up of 8 to 10 amino acids, which have a disruptive physiochemical effect resulting in alternation of membrane permeability in bacteria (52). In addition, type II NADH-quinone oxidoreductases, which are integral part of electron transport chain, has also been shown to be a secondary target sites of cationic peptides (53). Polymyxins have been administered for urinary tract infection, pneumonia, bacteremia, postoperative wound infections, abscesses, osteomyelitis (when given as an irrigation), and endocarditis (52).

Overall, we presented here a methodology that has been designed to challenge paradigms regarding metabolic dormancy in persisters, shed light on the often-overlooked metabolic processes of persister cells, develop a screening approach to identify metabolic inhibitors among a small library with FDA approved compounds, and integrate all proposed work to accelerate development of antipersister adjuvant therapies. Given that the cytotoxicity, cell permeability, solubility, and safety properties of FDA compounds have been well studied and documented during their preclinical and clinical research phases, discovering antipersister drugs among such libraries will have an enormous impact, because it will identify potential therapeutics that do not require the long laborious FDA approval process. Our preliminary studies have already identified a subset of drugs that can eliminate persisters even in stationary phase cultures, which represent notoriously challenging conditions for the elimination of persisters.

## MATERIALS AND METHODS

### Bacterial Strains and Plasmids

*Escherichia coli* MG1655 wild-type (WT) and MO strains as well as pQE-80L plasmids harboring genes encoding degradable (ssrA-tagged) green fluorescent protein (GFP) were obtained from Dr. Mark P Brynildsen at Princeton University. *Pseudomonas aeruginosa* PAO1 was a gift from Dr. Vincent Tam at the University of Houston. *E. coli* MO strain harbors a chromosomally integrated isopropyl β-D-1-thiogalactopyranoside (IPTG)-inducible *mCherry* expression cassette, which is used to monitor cell proliferation at single cell level (12, 14, 26). pQE-80L expression system has an IPTG-inducible synthetic *T5* promoter and a strong constitutive *LacI*^*q*^ promoter (with a point mutation) as a repressor, enabling us to tightly regulate the expression of *gfp* or *ssrA-gfp* (12). To directly measure protein degradation rates in stationary-phase cultures, we employed an assay using ssrA, a short peptide degradation tag with 11 amino acid residues that is linked to GFP to mark it for degradation by cellular proteases (27–30, 54). The overexpression of fluorescent proteins on *E. coli* persistence was shown to be insignificant (12–14, 26).

### Media, Chemicals and Culture Conditions

All chemicals were purchased from Fisher Scientific (Atlanta, GA), VWR International (Pittsburg, PA) or Sigma Aldrich (St. Louis, MO). Luria-Bertani (LB) liquid media, prepared from its components (5 g yeast extract, 10 g tryptone and 10 g sodium chloride in 1 L ultra-pure DI water), and Mueller-Hinton (MH) liquid media (21 g premixed MH in 1 L ultra-pure DI water) were used to grow *E. coli* and *P. aeruginosa*, respectively (12, 55, 56). LB agar media (40 g premixed LB agar in 1 L ultra-pure DI water) and MH agar media (38 g premixed MH agar in 1 L ultra-pure DI water) were used to enumerate the colony forming units (CFUs) of *E. coli* and *P. aeruginosa* strains, respectively. Phosphate Buffered Saline (PBS) solution was used to wash the cells to remove the chemicals and antibiotics before plating them on agar media. Concentrations of 5 µg/mL ofloxacin and 200 µg/ml ampicillin were used for persister assays (12). MIC ranges for *E. coli* MG1655 were found to be 3.125-6.25 µg/ml for ampicillin and 0.039-0.078 µg /ml for ofloxacin by using a method based on serial 2-fold dilutions of antibiotics in 2 ml LB media in 14 ml test tubes (13). MIC range for *P. aeruginosa* (PA01) were found to be 0.3125-0.625 µg/ml for ofloxacin. For selection and retention of plasmids in bacteria, 50 µg/ml kanamycin was added in culture media (12). To induce fluorescent protein expression, 1 mM IPTG was used (12). Primary drug screening was performed using Phenotype MicroArrays (PM11-20) in 96-well plate formats, containing various chemicals including FDA approved compounds (Biolog Inc., Hayward, CA). Eleven chemicals, identified as initial hits, were purchased separately for further investigation: amitriptyline hydrochloride (Fisher catalog#50-144-4347), trifluoperazine hydrochloride (Fisher catalog#T28495G), thioridazine hydrochloride (Fisher catalog#30-705-0), chlorpromazine hydrochloride (Fisher catalog#C24815G), CCCP (Fisher catalog# 04-525-00), protamine sulfate (Fisher catalog# AAJ6292609), promethazine hydrochloride (Fisher catalog#P2029100G), dodecyltrimethyl ammonium bromide (Fisher catalog# D146825G), triclosan (Fisher catalog# 64-795-01GM), polymyxin B Sulfate (Fisher catalog# 52-915-GM) and poly-L-lysine hydrochloride (VWR catalog# IC15269080). All chemicals were dissolved in ultra-pure DI water followed by filter-sterilization, except for CCCP and triclosan which were dissolved in DMSO. All LB and MH media were sterilized by autoclaving. Overnight precultures were prepared in 14-ml falcon tube containing 2 ml LB broth inoculated from a 25% glycerol (−80 °C) cells stock and grown for 24 h at 37 °C with shaking (250 rpm). Overnight pre-cultures were diluted in fresh 2 ml media in 14-ml test tubes or 25 ml media in 250-ml baffled flasks for the subsequent assays as describe below. Cells cultured in the presence of the solvent (DI water or DMSO) served as controls when the cultures were treated with chemical inhibitors.

### Cell Growth and Persister Assays

Overnight pre-cultures were diluted 1000-fold in 2 ml fresh LB media in test tubes and grown at 37 °C with shaking. Cell growth was monitored up to 24 hours by measuring optical density at 600 nm wavelength (OD_600_) with a plate reader (Varioskan LUX Multimode Microplate Reader, ThermoFisher, Waltham, MA) for selected time points. Cells were treated with chemicals at early-stationary (t=5 h) when required. At late-stationary phase (t=24 h), cells were diluted in 2 ml fresh media (yielding ∼5×10^7^ cells/ml) with antibiotics (5 µg/ml ofloxacin or 200 µg/ml ampicillin) in test tubes and incubated at 37 °C with shaking. At designated time points (t=0, 1, 2, 3, 4, 5 and 6 h), 100 µl samples were collected and washed with PBS to dilute the antibiotics to sub-MIC levels, followed by resuspension in 100 μl of PBS. Ten microliters of the cell suspension were serially diluted and plated on LB agar media to enumerate CFUs. The remaining 90 µl cell suspensions were also plated to increase the limit of detection. The agar plates were incubated at 37 °C for 16 h, which was found to be sufficient for *E. coli* colony formation (data not shown). The procedures described above were repeated using 250 ml-baffled flasks with 25 ml media to determine the effects of culture volume, aeration and mixing on cell growth and persistence. We did not see significant differences between the results of flask and test tube experiments (data not shown).

### Redox Sensor Green Dye Staining

Stationary-phase reductase and electron transport chain activities were measured with Redox Sensor Green (RSG) dye (ThermoFisher, catalog# B34954) according to manufacturer’s instructors. Cells at late-stationary phase (t=24 h) were diluted 100-fold in 1 ml PBS in flow cytometry tubes (5 ml round bottom falcon tubes, size: 12×75 mm) and stained with RSG at 1 μM concentration. For negative controls, CCCP (10 μM) was added in the cell suspensions 5 minutes before RSG staining to disrupt membrane electron transport. Mid-exponential-phase cells were used as positive controls (26, 43). Samples were incubated at 37 °C for 10 minutes before analyzing with a flow cytometer (NovoCyte Flow Cytometer, NovoCyte 3000RYB, ACEA Biosciences Inc., San Diego, CA). Forward and side scatter parameters of unstained controls were used to gate the cell populations on flow diagrams. Cells were excited at 488 nm with solid-state laser, and green fluorescence was collected with a 530/30 bandpass filter. To analyze the effect of chemical inhibitors (e.g., chlorpromazine) on stationary-phase-cell metabolism, cells at early-stationary phase (t=5 h) were treated with the chemicals at indicated concentrations, and RSG staining was performed at t=24 h as described above.

### Monitoring Cell Division and Quantifying VBNC Cells

To monitor cell division and quantify non-growing cell subpopulations, inducible fluorescent protein (mCherry) expression systems were used. Overnight pre-cultures of *E. coli* MO strain were diluted 1000-fold in 2 ml LB media with 1 mM IPTG (to induce *mCherry*) in test tubes and grown as described. We previously showed that mCherry expression cassette or overexpressing mCherry did not affect the *E. coli* persistence (12, 14, 26). If necessary, cells at early-stationary phase (t=5 h) were treated with chemical inhibitors (e.g., chlorpromazine) at indicated concentrations. At t=24h, mCherry-positive cells were collected, washed twice with PBS to remove the IPTG, resuspended in 25 ml fresh LB media without inducer in 250 ml baffled flasks and cultured at 37 °C with shaking. At designated time points (t= 0, 1, 2, and 2.5 h), cells were collected, washed and resuspended in PBS to measure their fluorescent protein content with a flow cytometer. When necessary, cells were further diluted in PBS to reach a desired cell density for the flow analysis (10^6^-10^7^ per ml). Cell division was monitored by measuring the dilution rate of fluorescent protein at single cell level. At t=0 h, all cells exhibited a high level of red fluorescence, which declined as the cells divided, except in a small subpopulation whose fluorescence remained constant due to the lack of division (t=2.5 h). Given that ampicillin only targets the proliferating cells, the cultures were further challenged with ampicillin (200 µg/ml) to quantify VBNC and persister cells in non-growing cell subpopulations. Using LIVE/DEAD staining, FACS and clonogenic survival assays, we previously showed that antibiotic sensitive cells were rapidly lysed by ampicillin while VBNC and persister cells remained intact (14). The intact cells were quantified using the volumetric-based cell counting feature of the NovoCyte Flow Cytometer. Persisters were quantified by enumerating the CFUs after plating the ampicillin treated cultures as described above. Intact cells that did not form colonies on standard medium were classified as VBNC cells (14, 21–25). All samples were assayed with lasers emitting at 561 nm and red fluorescence was collected by 615/20 nm bandpass filter.

### Fluorescent Protein Degradation Assay

Overnight pre-cultures of *E. coli* MG1655 harboring pQE-80L-*ssrA*-*gfp* were inoculated (1:1000) in 2 ml LB in test tubes, grown in the presence of IPTG (to induce *ssrA* tagged *gfp*) until the early stationary phase (t=5 h). Then, the cells were washed to remove the inducer, resuspended in filter-sterilized 2 ml spent medium (obtained from cultures grown under identical conditions without the inducer) and cultured in test tubes at 37 °C with shaking. When necessary, cell suspensions were treated with chemical inhibitors. At designated time points, 200 µl samples were collected to measure their GFP levels with a plate reader. Excitation and emission wavelengths for GFP detection was 485 nm and 511 nm, respectively.

### Chemical Screening

Early-stationary-phase cells expressing ssrA-tagged GFP (grown in 25 ml LB with IPTG in 250 ml baffled flasks) were washed, resuspended in spent medium (without inducer), transferred to 96-well PM plates (100 µl per well) containing the chemical library, covered with sterile, oxygen-permeable sealing membranes, and cultured in a shaker at 37 °C and 250 rpm. GFP levels were monitored for 4 h (which was found to be sufficient) using a plate reader, with cells cultured in the presence of the solvent serving as the negative controls, and those with chlorpromazine as a positive control. GFP measurements taken at 4 h were normalized to those taken at 0 h to eliminate any variations in initial cell concentrations. Z-score method, calculated from the mean and the standard deviation of all measurements within the plate (31) was used to determine initial hits:

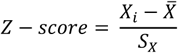

where *X*_*i*_ is the measurement (normalized) of the *i*^*th*^ compound, 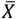 and *S*_*X*_ are the mean and the standard deviation of all measurements. An absolute Z-score of ≥2, which correlates to a P-value of 0.045 (32), was assumed to be the threshold for hit detection. We note that each plate contains four different concentrations for each compound (information on these concentrations not disclosed by the company). Z-scores were calculated for each concentration set. The initial hits were selected among the chemicals that successfully inhibited GFP degradation (Z-score≥2) with at least two different concentrations.

Assay validation was evaluated by Z-factor calculated from the mean and standard deviation values of the positive (p) and the negative (n) control plates, as follows:

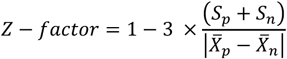

A Z-factor between 0.5 and 1.0 indicates that the proposed assay is robust and reliable (57).

### Validating the Selected Chemicals

To fully assess their utility and effectiveness, the selected chemical hits were analyzed at various concentrations with the aforementioned assays. Overnight pre-cultures of *E. coli* strains (WT, MO or cells expressing *ssrA* tagged *gfp*) were inoculated (1:1000) in 2 ml LB (IPTG added for the cells harboring inducible fluorescent proteins) in test tubes and cultured as described. Cells at t=5 h were treated with chemicals at indicated concentrations. Fluorescent protein degradation assays throughout the stationary phase after the treatments were performed for the cultures of *E. coli* cells expressing SsrA tagged GFP; persister and cell survival assays at late-stationary phase (t=24 h) were performed for WT cultures; and finally, cell division assays at late-stationary phase were performed for the *E. coli* MO cultures.

### *Pseudomonas aeruginosa* Persister Assay

Overnight pre-cultures of *Pseudomonas aeruginosa* (PA01) were inoculated (1:1000) in 2 ml MH broth in test tubes and cultured as described before. At early stationary phase (t=5 h), cells were treated with chemicals at indicated concentrations. At t=24 h, cells were washed to remove chemicals and inoculated (1:100) in 1 ml MH broth followed by ofloxacin (5 µg/ml) treatment. At t=0 (before ofloxacin treatments), ten microliter cell suspensions were serially diluted and spotted on MH agar media to enumerate initial CFUs, which enables us to assess the impacts of chemical treatments on *P. aeruginosa* (PA01) cell viability. To enumerate persister levels at t=6 h, ofloxacin treated cultures were washed, serially diluted and plated on MH agar media to incubate 20 h at 37 °C. Twenty-hour incubation was found to be sufficient for *P. aeruginosa* (PA01) colony formation (data not shown).

### Statistical analysis

Two tailed t-test with unequal variances was used to evaluate the statistical significance, where P<0.05 (55). At least three independent biological replicate was performed for all experiments. All data points on linear-scale graphs indicate mean value ± standard error; however, for logarithmic-scale graphs, standard deviations were used to better represent the error bars.

## ACKNOWLEDGEMENTS

We would like to thank Dr. Aslan Massahi for providing assistance in persister assays.

## Author Contributions

S.G.M, T.H, A.S., P.K. and M.A.O. conceived and designed the study. S.G.M, T.H, A.S. and P.K. performed the experiments. S.G.M., T.H, A.S., P.K. and M.A.O. analyzed the data and wrote the paper.

## Data and materials availability

Data provided in this paper including supplementary materials are sufficient to assess the findings of this paper. Additional data of this paper can be obtained upon request.

