## Supplemental File for "Therapeutically exploring persister metabolism in bacteria"

### SUPPLEMENTARY MATERIALS

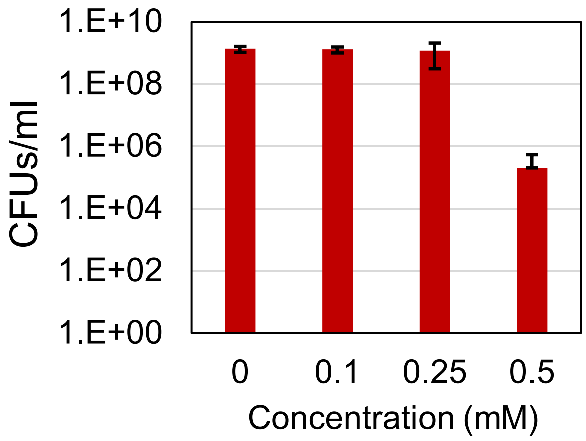


**Fig. S1. Chlorpromazine (CPZ) treatment throughout the stationary phase.** Cells at early-stationary phase (t=5 h) were treated with CPZ at indicated concentrations, and then, cells at late stationary phase (t=24 h) were washed to remove the chemicals and plated on agar media to assess the effects of CPZ treatments on cell viability.

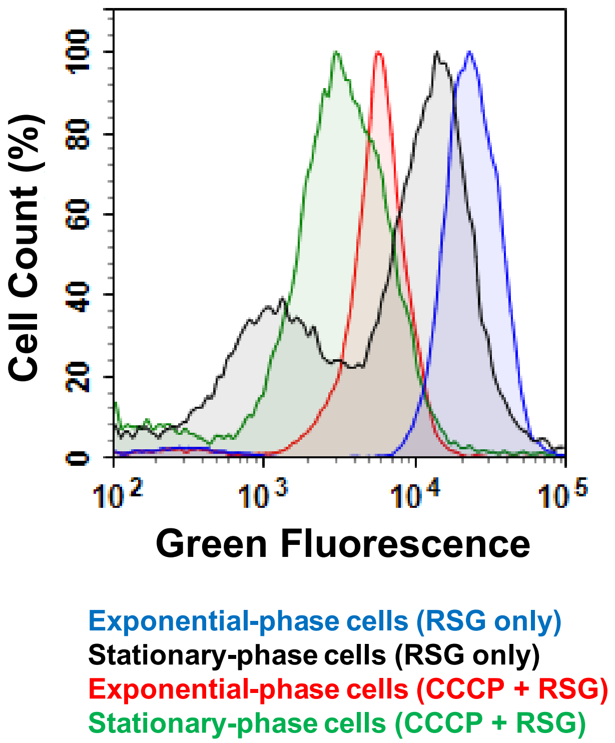


**Fig. S2. RSG is an indicator of bacterial reductase activity.** Cells at mid-exponential phase (t=3h) and late-stationary phase (t=24h) were transferred to PBS and stained with RSG. For controls, cells were treated with a metabolic inhibitor, CCCP, as described in the manufacturer's protocol.

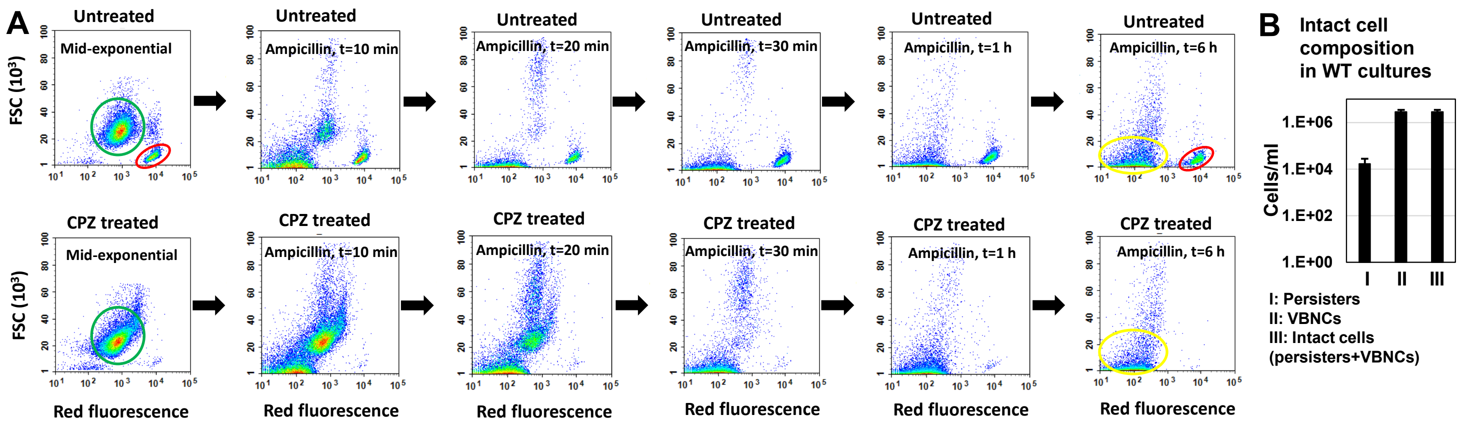

**Fig. S3. Chlorpromazine (CPZ) treatment throughout the stationary phase reduced VBNC cell formation.** Overnight pre-cultures of MO cells (harboring an inducible mCherry expression system, i.e., pQE-80L-*gfp*) were diluted 1000 fold in 2 ml LB in test tubes in the presence of inducer (1 mM IPTG) and cultured. Cells at early stationary phase (t=5h) were treated CPZ (0.25 mM) or left untreated with washed. At t=24h, cells were washed to remove the inducer and transferred to fresh media without IPTG to monitor the cell growth. All cells exhibited high red fluorescence at t=0 h (Fig. 1F, main text), and the red fluorescence signal declined as the cells divided (growing cells highlighted with a green circle), except for small subpopulations in which the fluorescence signal remained constant due to the lack of cell division (non-growing cells highlighted with a red circle). We note that the 1st column in Panel A corresponds to the last row of Fig. 1F in the main text. VBNC cell levels were determined with the ampicillin-treatment approach as described previously. When the cells at mid-exponential phase were treated with ampicillin, the growing cell subpopulations were lysed (debris highlighted with a yellow circle), however non-growing cells remained intact. Only a small fraction of intact cells (i.e., persister cells) colonized (panel B). The majority of intact cells were detected as VBNC cells (panel B). Non-growing cells were not detectable in CPZ treated cultures (panel A). Note that a representative biological replicate is shown here. All 3 biological replicates consistently resulted in similar trends.

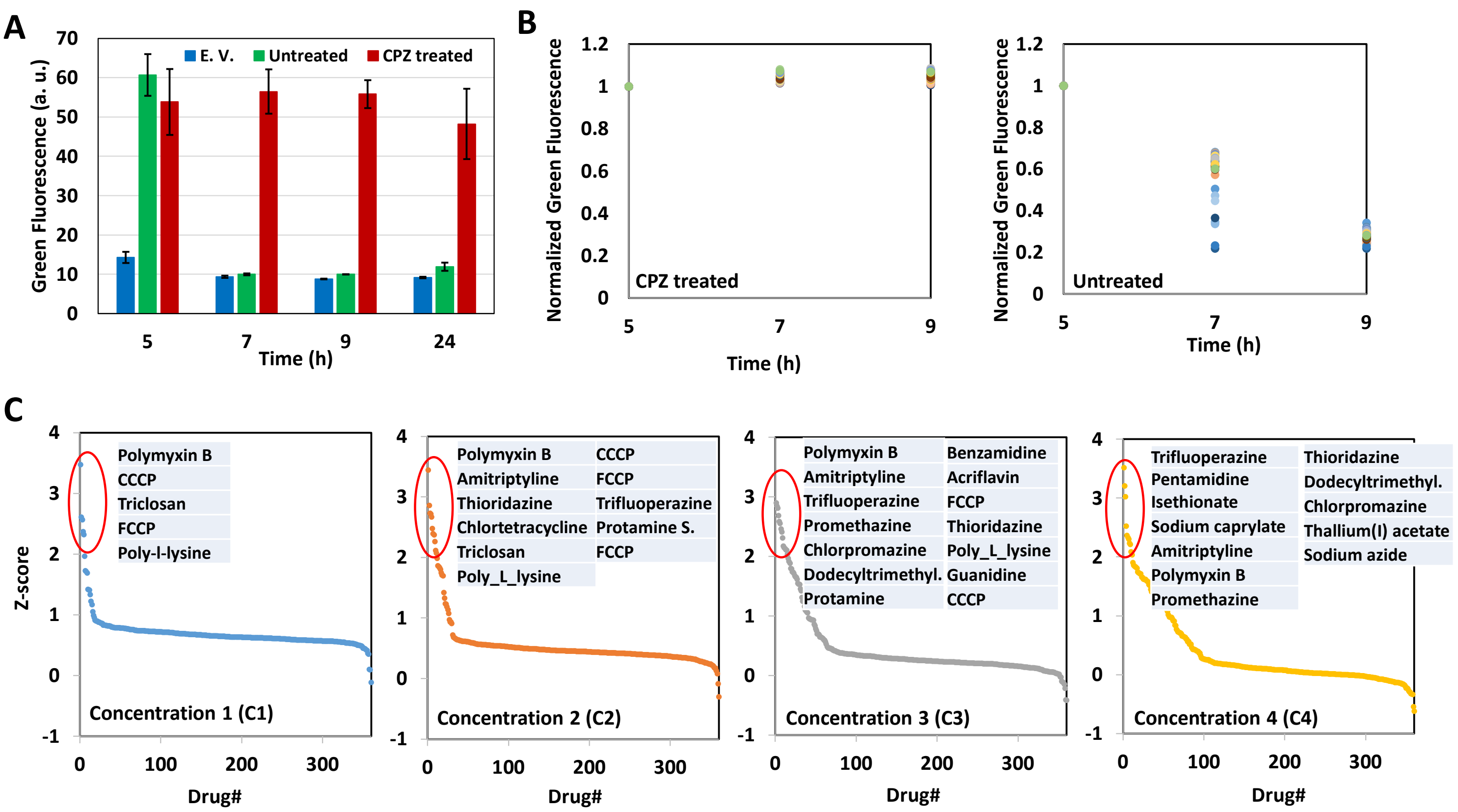


**Fig. S4. Developing a chemical screening approach.** (A) Stationary-phase GFP degradation was inhibited with CPZ treatment (0.25 mM). Cells expressing pQE-80L*gfp-ssrA* were grown to stationary phase (t=5h) in the presence of IPTG (inducer) and then re-suspended in a filter-sterilized spent medium (without inducer and obtained from the cultures grown under identical conditions) and immediately treated with CPZ to inhibit cell metabolism and protein degradation. Untreated cell cultures (no CPZ treatment) served as negative control. Background fluorescence was determined using cells with empty vectors (E.V.). (B) Stationary-phase cells expressing ssrA-tagged GFP were re-suspended in spent medium, without inducer, transferred to 96-well plates treated with CPZ or left untreated, and cultured in a shaker. GFP measurements were taken for 4 h and normalized to those taken at 0 h. (C) Stationary-phase cells expressing ssrA-tagged GFP were re-suspended in spent medium, without inducer, transferred to 96-well PM plates containing the chemical library, and cultured in a shaker for 4h. GFP measurements taken at 4 h were normalized to those taken at 0 h. The Z-scores calculated for the chemical compounds at four different concentrations ($C_4 > C_3 > C_2 > C_1$). Chemicals with Z-scores >2 were tabulated for each concentration set. Eleven hits were selected among the chemicals that successfully inhibited GFP degradation (Z-score≥2) with at least two different concentrations (Fig. 2C, main text).

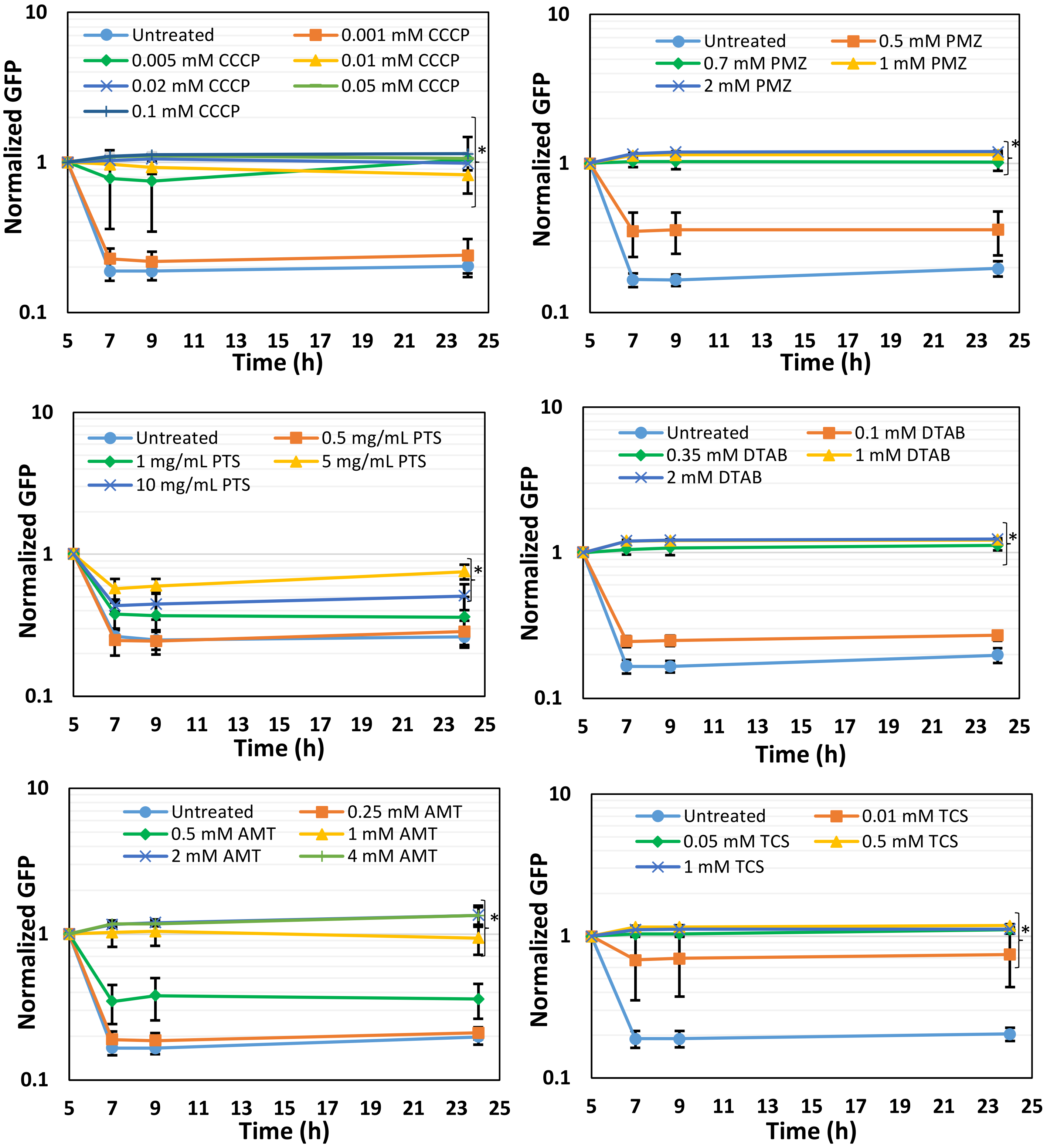


**Fig. S5. Determining inhibitory concentrations for GFP degradation.** Cells were treated with hit drugs at early stationary phase (t=5h) at indicated concentrations, and then, GFP measurements were performed at indicated time points. Eleven hits, selected based on the Z-scores analysis (Fig.

2C and Fig. S4), were analyzed. Six chemicals were highlighted in this figure; the rest of the chemicals were highlighted in Fig. 2D in the main text. TDZ: Thioridazine; PLL: Poly-L-lysine; PMXB: Polymyxin B; TFP: Trifluoperazine; CCCP: Carbonyl cyanide m-chlorophenyl hydrazone; PMZ: Promethazine; PTS: Protamine Sulfate; AMT: Amitriptyline; DTAB: Dodecyltrimethylammonium bromide; TCS: Triclosan; (N=3).

*: Statistical significance between drug-treated vs. untreated cultures at last three time points ($P<0.05$, two-tailed t-tests with unequal variances).

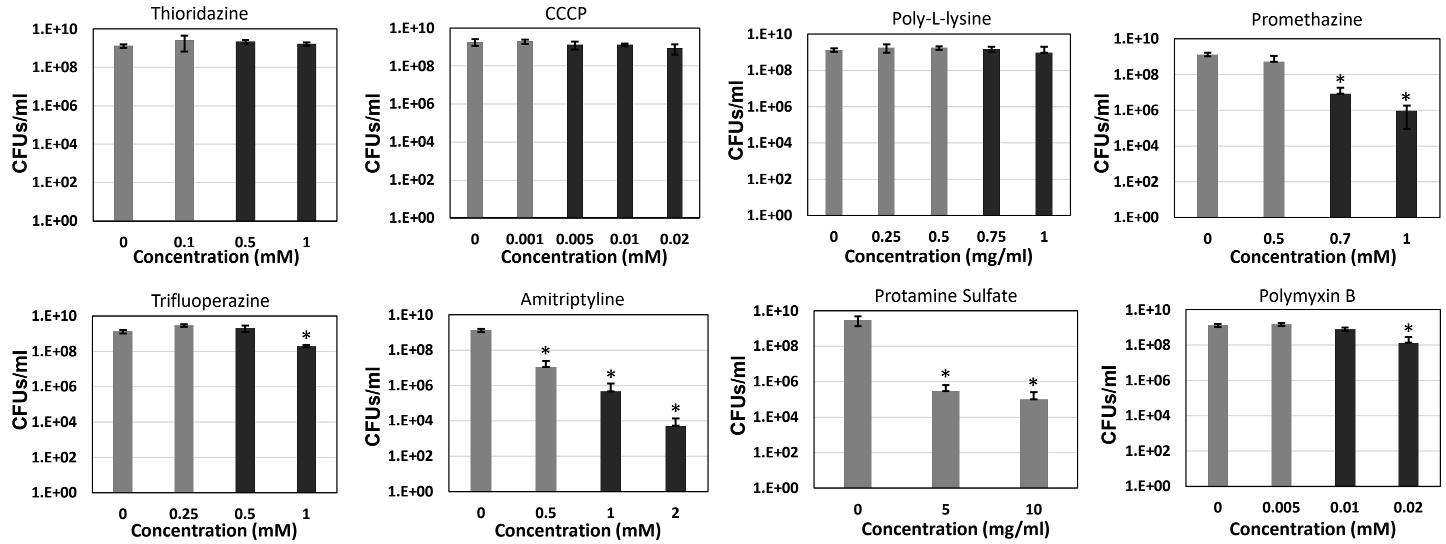


**Fig. S6. The effects of chemical hits on cell viability.** Cells at early-stationary phase were treated with chemical hits at various concentrations, and then, cells at late-stationary phase were washed to remove the chemicals and plated on agar media for CFU measurements. Among the chemicals tested, CCCP, Polymyxin B, Poly-L-lysine, Thioridazine, Trifluoperazine did not affect the cell viability within a wide range of concentrations tested. Triclosan and Dodecyltrimethyl ammonium bromide significantly reduced the stationary phase cell survival at concentration ranges that inhibit GFP degradation (data not shown). Black columns: concentrations that inhibit GFP degradation. Grey columns: concentrations that do not inhibit GFP degradation. * indicates a significant reduction in treated groups compared to untreated controls ($P<0.05$, two-tailed t-tests with unequal variances), (N=3).

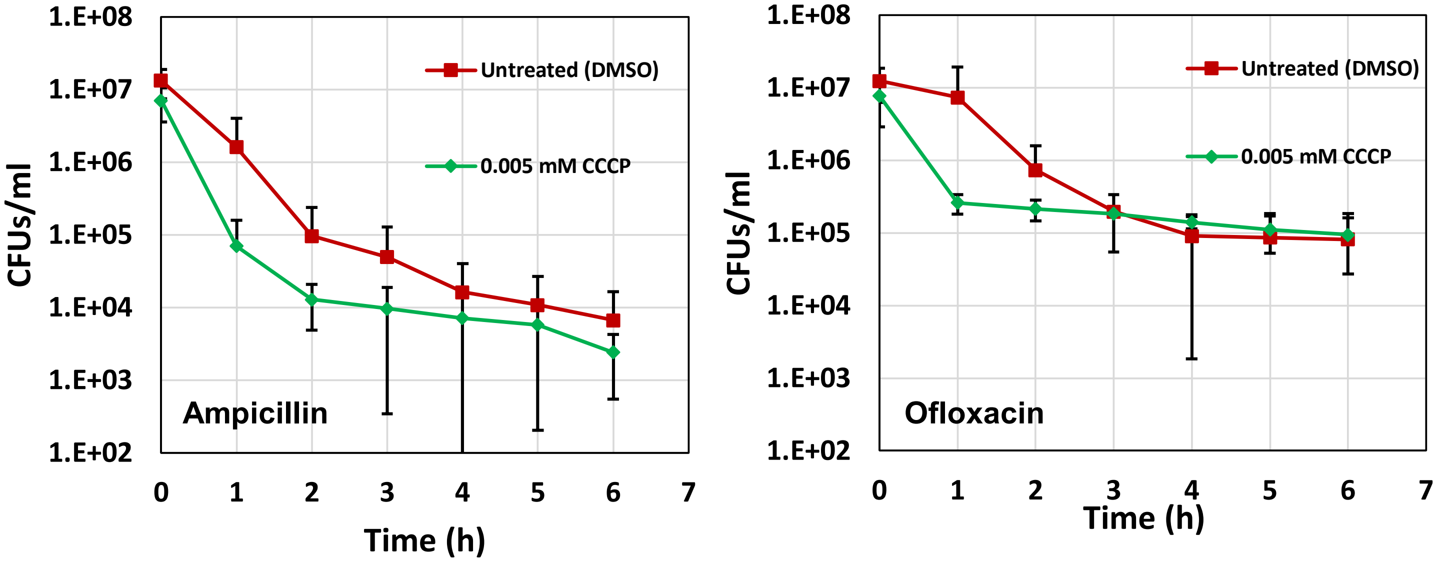


**Fig. S7. CCCP pre-treatment did not reduce *E. coli* persistence.** Cells at early-stationary phase were treated with the five chemicals (CCCP, Polymyxin B, Poly-L-lysine, Thioridazine, and Trifluoperazine, highlighted in Fig. S6) at concentrations that inhibit GFP degradation without affecting the stationary-phase-cell survival. Then, cells at late stationary phase were washed to remove the chemicals, transferred to fresh media and treated with ofloxacin and ampicillin. CCCP data was provided here; the data for the rest of the chemicals were provided in the main text (Fig. 2). We observed large standard deviations in the persister levels of CCCP treated cultures (N=3).

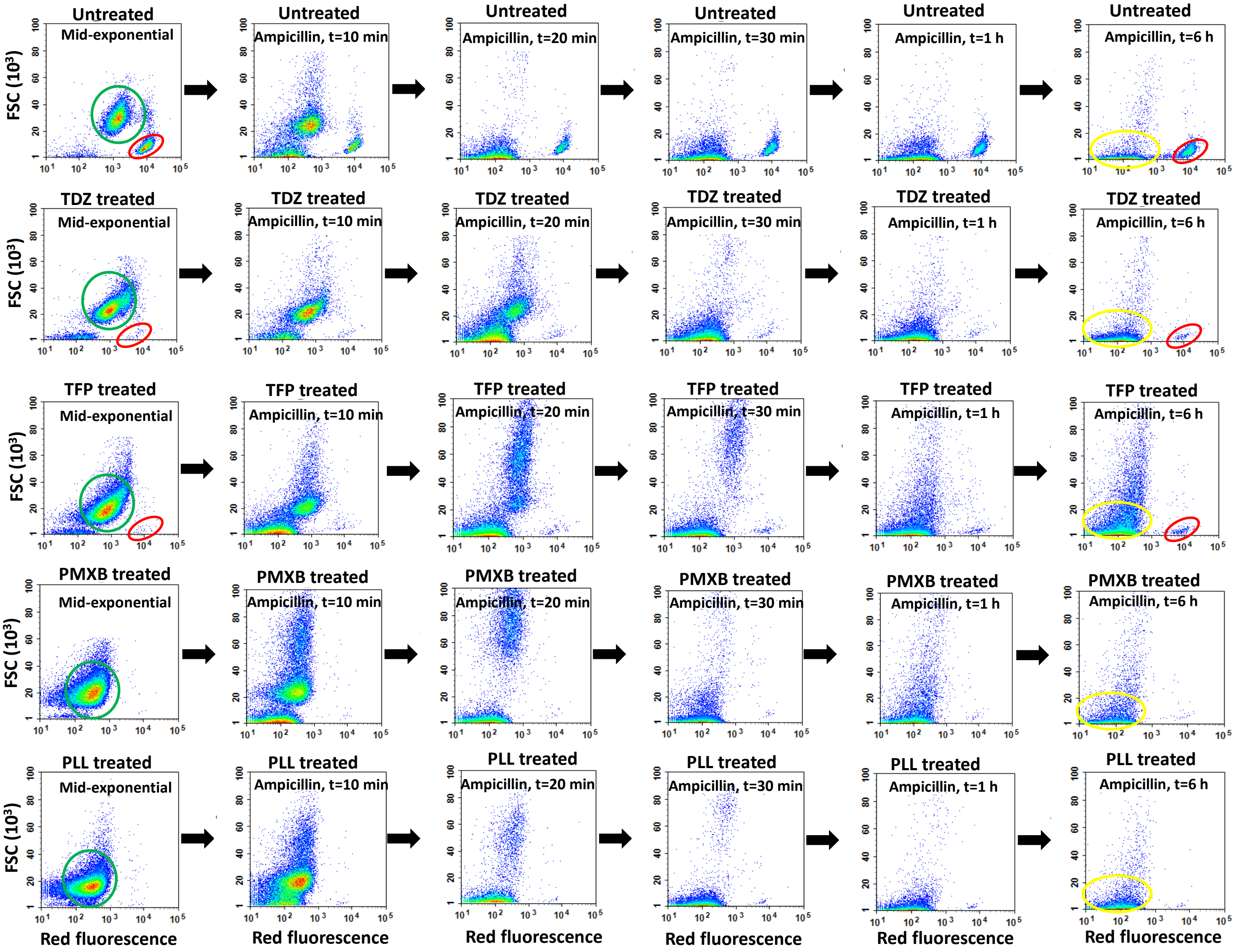


**Fig. S8. Treatment of stationary-phase cells with the identified chemicals reduced VBNC cell formation.** Growing cells (green circle), non-growing intact cells (red circle) and debris (yellow circle) were determined as described previously. Note that a representative biological replicate is shown here. All 3 biological replicates consistently resulted in similar trends. Drug concentrations: 0.5 mM Thioridazine (TDZ); 0.5 mM Trifluoperazine (TFP); 0.01 mM Polymyxin B (PMXB); 0.75 mg/ml Poly-L-lysine (PLL).

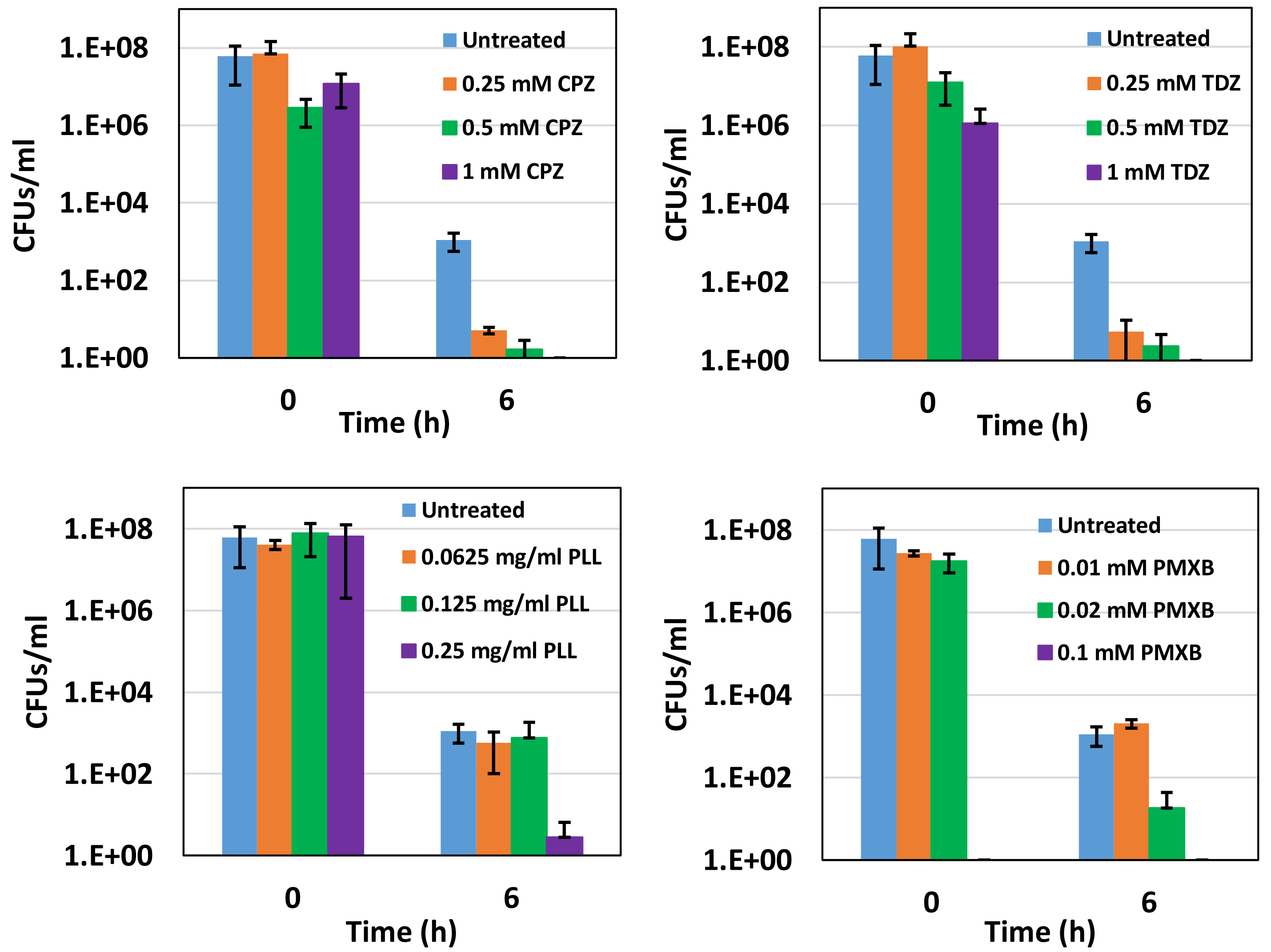


**Fig. S9. Persister levels in *P. aeruginosa* cultures treated with the chemical hits.** Early-stationary-phase cells (t=5h) were treated with the selected drugs or left untreated (control); cells in late stationary phase were then washed to remove inhibitors and re-suspended in fresh media with ofloxacin (effective for *P. aeruginosa*) for persister assays. Cells were plated for CFU enumeration before and after the ofloxacin treatments to assess the effects of drugs on *P. aeruginosa* cell viability and persistence, respectively, (N=3).
